# Advanced Retinal Imaging and Ocular Parameters of the Rhesus Macaque Eye

**DOI:** 10.1101/2020.09.12.294165

**Authors:** Kira H. Lin, Tu Tran, Soohyun Kim, Sangwan Park, J. Timothy Stout, Rui Chen, Jeffrey Rogers, Glenn Yiu, Sara Thomasy, Ala Moshiri

**Author notes:** Corresponding author: Ala Moshiri M.D. Ph.D., Department of Ophthalmology and Vision Science, School of Medicine, University of California at Davis, Eye Center, 4860 Y. Street, Suite 2400, Sacramento, CA 95817.

## Abstract

**Purpose:** To determine the normal ocular biometric and perform advanced retinal imaging and functional assessment of a non-human primate used commonly in scientific research, the rhesus macaque.

**Methods:** We performed ocular phenotyping on rhesus macaques at the California National Primate Research Center. This consisted of anterior and posterior segment eye examination by ophthalmologists, advanced retinal imaging, and functional retinal electrophysiology.

**Results:** Full eye exams were performed on 142 animals consisting of pupillary light reflex, tonometry, external exam and photography, anterior slit lamp examination, and posterior segment examination by indirect ophthalmoscopy. Ages of the rhesus macaques ranged from 0.7 to 29 years (mean=16.4 years, stdev=7.5 years). Anterior segment measurements such as intraocular pressure (n=142), corneal thickness (n=84), lens thickness (n=114), and axial length (n=114) were acquired. Advanced retinal imaging in the form of fundus photography (n=78), optical coherence tomography (n=60), and quantitative autofluorescence (n=44) were obtained. Electroretinography (n=75) was used to assay retinal function. Quantitative analyses of macular structure, retinal layer segmentation, and rod and cone photoreceptor electrical responses are reported. Quantitative assessments were made and variations between genders and age groups were analyzed to compare with established sex and age-related changes in human eyes.

**Conclusions:** The rhesus macaque has ocular structure and function very similar to that of the human eye. Age-related ocular changes between rhesus and humans are similar. In particular, macular structure and function are very similar to humans making this species particularly useful for the study of macular biology and development of therapies for inherited and age-related macular degenerations as well as cone photoreceptor disorders.

## Introduction

Vision science research has relied on small animal models, particularly rodents, for the bulk of eye research in the last several decades. Mice in particular are a commonly used animal model of human eye disease since they are easy to breed, are small in size, have short gestational periods, and develop to adulthood relatively quickly. Furthermore, the mouse genome is similar to the human genome and is routinely manipulated producing many genotype-phenotype correlations that are parallel between species. However, mouse eyes are comparatively small, and the relative size of ocular structures are substantially different from people. The mouse eye has a large lens and small vitreous in comparison to the human eye, and it lacks a high acuity area, also referred to as the macula. When studies require larger eyes, rabbits, pigs, and dogs serve as useful models. Rabbits have been used particularly for ocular toxicology studies.^1^ Pigs have been used as models of ocular and retinal surgery.^2–4^ Select forms of canine models of inherited retinal disease have also been characterized.^5,6^ However, none of these species have a macula, which represents a major limitation for studying the most common forms of human retinal blindness. Similarly, models of corneal disease and glaucoma in these species are limited in number and similarity to human ophthalmic disease. Furthermore, the overall shape and structure of the primate eye has unique adaptations not found in other mammals. Therefore, precise modeling of all forms of human blindness requires the use of non-human primates (NHPs). One of the commonly used NHPs used for biological research are the rhesus macaque (*Macaca mulatta*).

Upon inspection, the rhesus macaque eye and the human eye bear a tremendous amount of anatomical similarity. The similarities are so striking that images of the NHP retina can be difficult to distinguish from the human even for ophthalmologists. In general, the cornea, iris, lens, and vitreous also appear grossly identical between NHPs and human patients.^7^ These commonalities make NHPs an attractive model for translational studies to develop new therapies for virtually all forms of human eye disease. The NHP eye, in particular, has a macula that is very similar to the human counterpart. Given that many important forms of blindness involve the macula, and that this specialized structure does not exist in rodents or even in most large animal models, NHP research is particularly well suited to the study and development of macular diseases and cone photoreceptor disorders.^8,9^ In order to pursue translational studies in NHPs, it is necessary to establish norms of ocular examination and structure, as well as retinal function and anatomy.

The purpose of this study was to develop a normative database of ocular structure and function based on complete eye examination, ocular measurements, advanced retinal imaging, and electroretinography. All data were derived from normal animals of various ages as part of a prospective screening process to identify animals with inherited retinal disease.^9^ We hypothesized that the anatomic imaging and electroretinogram (ERG) in the NHP eye would effectively mirror human retinal norms with possible species-specific differences. We collected intraocular pressure (IOP), central corneal thickness, lens thickness, and axial length. We measured anatomical parameters of the macula from fundus photography, retinal layer thicknesses on optical coherence tomography (OCT), and retinal pigment epithelial lipofuscin using quantitative autofluorescence (qAF). We analyzed OCT images for retinal structure and anatomy focusing on the macular architecture. Quantitative autofluorescence has potential in human ophthalmology practice,^10–12^ and given the utility of NHP eye research, we provide qAF norms for comparison with human clinical data. All ophthalmic data were analyzed with regard to sex and age to identify sex-specific differences and findings associated with aging. Known associations in humans such as increased IOP measurements correlated with central corneal thickness were tested. We also analyzed whether or not IOP, axial length, prevalence of punctate macular lesions, thickness of retinal layers on OCT, and qAF vary with age. Our investigations reveal the eye of the rhesus macaque is structurally and functionally similar to the human eye in essentially every measure and undergoes changes over the lifespan of the animal which are relevant to many forms of human age-related eye diseases.

## Methods

### Animals

All of the animals in this study were rhesus macaques (*Macaca mulatta*) born and maintained at the California National Primate Research Center (CNPRC). The CNPRC is accredited by the Association for Assessment and Accreditation of Laboratory Animal Care (AAALAC) International. Guidelines of the Association for Research in Vision and Ophthalmology Statement for the Use of Animals in Ophthalmic and Vision Research were followed. All aspects of this study were in accordance with the National Institutes of Health (NIH) Guide for the Care and Use of Laboratory Animals. Phenotyping and ophthalmic examinations were performed according to an animal protocol approved by the UC Davis Institutional Animal Care and Use Committee.

### Ophthalmic Phenotyping

Intramuscular injection of ketamine hydrochloride and dexmedetomidine were administered for sedation. Animals were monitored by a trained technician and a veterinarian at all times.

Ophthalmic examination included measurement of intraocular pressure using rebound tonometry (Icare TA01i, Finland), pupillary light reflex testing, external and portable slit lamp examination, as well as dilated (Tropicamide 1%, Phenylephrine 2.5%, Cyclopentolate 1%) indirect ophthalmoscopy. Axial length and lens thickness were measured using a Sonomed Pacscan Plus (Escalon, Wayne, PA, United States). Corneal thickness was measured using a DGH Pachette 4 (DGH Technology Inc, Exton, PA, United States).

### Ophthalmic Imaging

The CF-1 Retinal Camera with a 50° wide angle lens (Canon, Tokyo, Japan) was used to obtain color and red-free fundus photographs. Spectral-domain optical coherence tomography (SD-OCT) with confocal scanning laser ophthalmoscopy (cSLO) was performed (Spectralis^®^ HRA+OCT, Heidelberg, Germany). The Heidelberg eye tracking Automatic Real-Time (ART) software was set at 25 scans for each B-scan. A 30-degree horizontal high-resolution raster scan centered on the fovea was obtained using a corneal curvature (K) value of 6.5 mm radius. Focal measurements of retinal layer thickness measurements were performed using the caliper measurement tool in ImageJ (National Institutes of Health, Bethesda, MD, United States) on a horizontal line scan through the foveal center. The Spectralis device was also used to obtain blue-peak fundus autofluorescence in a quantitative fashion as described below.

### Electroretinography

A dark and light adaption full-field ERG (ffERG) containing six different tests was performed on each eye following a 30 minute dark adaptation. ERG-Jet electrodes (item #95-011) were coupled with the RETeval instrument (LKC Technologies, Gaithersburg, MD, United States). A standard flash electroretinogram was performed according to the approved protocol of the International Society for Clinical Electrophysiology of Vision (ISCEV). There were four dark adapted tests (0.01 cd*s/m^2^, 3.0 cd*s/m^2^, 10.0 cd*s/m^2^, and oscillatory potentials 3.0 cd*s/m^2^). After 10 minutes of light adaptation, two additional tests were performed (3.0 cd*s/m^2^ and flicker 3.0 cd*s/m^2^). Both time (ms) and amplitude (μV) were obtained for each test on each eye. Single flash tests measured an a-wave and b-wave. Oscillatory potentials measured five wave points and a sum. In the photopic flicker test, the first wave point is reported. Left and right eye measurements were combined and averaged for each test per primate. Measurements were recorded and displayed using the manufacturer’s software. Recordings from ERG tests that were of poor quality or non-measurable were discarded.

### Fundus Measurements

Fundus images were obtained in both monochromatic red-free and color. All measurements were made using red-free images, except in rare instances when the quality of the color image was superior to the former. Measurements were originally measured in pixels and then converted to millimeters (mm) based on the conversion factor for a 50° fundus camera of 194.98 pixels/mm.^13^ Images were rotated so that the vertical diameter of the optic nerve was maximized, such that the superotemporal and inferotemporal macular arcades of blood vessels were roughly parallel to one another. The measurements for the two eyes of each animal were combined and averaged. Measurements included the largest arterial and venous diameters at their crossing with the edge of the optic disc and the arterial-venous ratio was calculated as the diameter of the artery over the diameter of the vein. The maximum vertical and horizontal diameters of the optic nerve were calculated. Macular parameters were measured including the distance from the foveal center to the optic nerve center, and the distance from the foveal center to the nerve edge. The area of the foveal avascular zone was estimated using both fundus photos and fluorescein angiography. Additionally, we measured the relative position of the foveal center with respect to the vertical diameter of the optic nerve head. The distance was measured from the nerve center point to the base of the optic nerve and was calculated as a percentage of the total optic nerve height.

### Optical Coherence Tomography (OCT)

Optical coherence tomographic images were captured during a sedated exam of each primate. For each eye, an OCT image was chosen where the fovea had the greatest depth. Measurements were taken using the linear tool in ImageJ. The manufacturer’s scale bars on the OCT image were used to calibrate the measurements. Thickness measurements of the retinal layers were taken at three locations: 1.5 mm temporal to the foveal center, at the foveal center, and 1.5 mm nasal to the foveal center. The layers measured included the nerve fiber layer (NFL), ganglion cell layer (GCL), inner plexiform layer (IPL), inner nuclear layer (INL), outer plexiform layer (OPL), outer nuclear layer (ONL), photoreceptor inner segments (IS), photoreceptor outer segments (OS), retinal pigmented epithelium (RPE), choriocapillaris (CC), outer choroid (OC), and total retinal thickness. The layers measured in the foveal center were the NFL, ONL, IS, OS, RPE, CC, and OC. Average thicknesses were calculated at each of these three locations for each eye of each primate.

### Quantitative Autofluorescence (qAF) Analysis

The methods for capturing qAF has been described by Delori et al^14^ and Greenberg et al.^15^ In brief, after SD-OCT was performed, the device was turned to qAF mode to capture 30° x 30° qAF images with excitation light of 488 nm and a long-pass barrier filter starting at 500 nm increasing to 80 nm, calibrated with an internal master fluorescence reference. Photobleaching was achieved by exposing the retina to 488 nm blue excitation light for 30 seconds. Images were captured from the central macula, with intensity adjusted using an internal fluorescence reference to enable quantification of autofluorescence (AF) and normalizing AF units given minor variations in laser power and detector sensitivity in between imaging sessions. For each eye, three series of 12 successive images were acquired in rapid succession; the mean image of the three sequences was computed using the manufacturer’s qAF software module. Then, for each mean image a Delori grid was centered on the fovea and expanded to the tangential edge of the optic nerve. The main measure for each image is the mean qAF8, which was acquired after selecting the middle eight segments to exclude vessels and other noise. Each image was then individually graded on a 0-3 scale (0 being unusable and 3 being the highest image quality) by two independent graders (K.H.L. and T.M.T.). The two independent graders reached consensus on each image’s final grade and averaged the qAF8 for each eye to arrive at the final qAF8 measurement for each eye. Only images graded 2 or above were evaluated and poor-quality images were excluded.

### Statistical Analysis

Descriptive statistics were used to summarize demographic and all ocular measurement data. Panel regression was used to determine statistical significance (*P* < 0.05) treating each eye as the unit of analysis linked by the primate to account for within-subject correlation, and each primate’s panel consisted of right eye and left eye measurements. Analyses were performed in Microsoft Excel (Microsoft Corporation, Redmond, WA, United States) and STATA 16 (StataCorp, College Station, TX, United States).

## Results

Data were collected from a total of 142 individuals. Ages of the rhesus macaques ranged from 0.7 to 29 years (mean = 16.4 years, stdev = 7.5 years). The age and gender distribution are shown in Figure 1.

**Figure 1.**
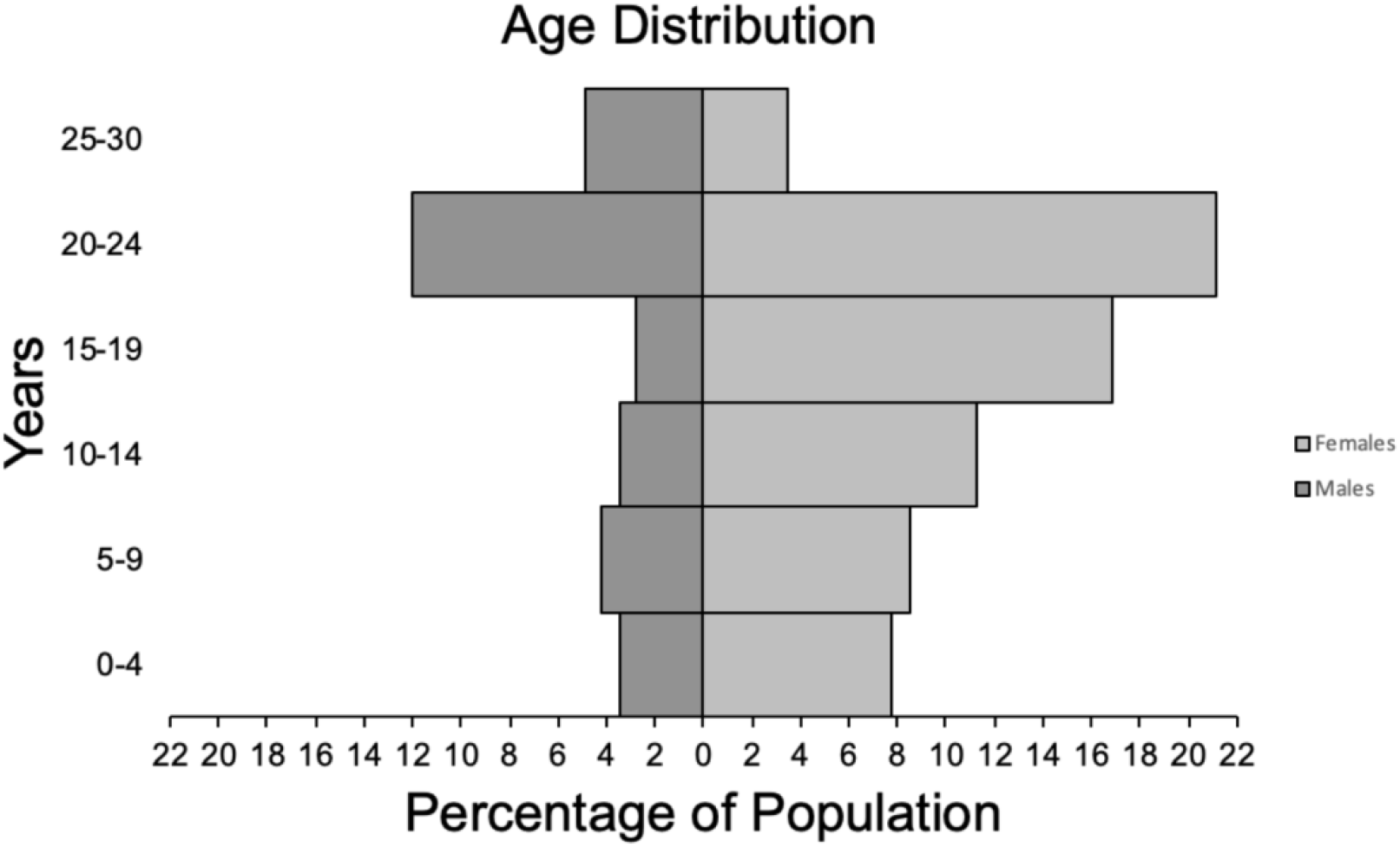
Age distribution of total rhesus macaques (n = 142) by sex (male and female). The mean was 16.4 years (stdev = 7.5 years). Median was 19.2 years. Macaques were separated into 6 different age groups: 0-4 years, 5-9 years, 10-14 years, 20-24 years, and 25-30 years. Age group 0-4 years old (M: 5, F: 11), age group 5-9 years old (M: 6, F: 12), age group 10-14 years old (M: 5, F: 16), age group 15-19 years old (M: 4, F: 24), age group 20-24 (M: 17, F: 30), and age group 25-30 (M: 7, F: 5).

Intraocular pressure (IOP) measurements were collected from a total of 142 individuals, 284 eyes (Figure 2). The mean ± SD IOP was 18 ± 4 mm Hg in all animals; 17 ± 4 mm Hg in the OD and 18 ± 5 mm Hg in the OS. The mean IOP in males was 19 ± 5 mm Hg and in females 17 ± 4 mm Hg. There was no statistical difference between right and left eyes (*P* = 0.369) or males and females (*P* = 0.088). We did find that IOP increased (Figure 2A) by 0.165 mm Hg per each year of age (*P* < 0.001).

**Figure 2.**
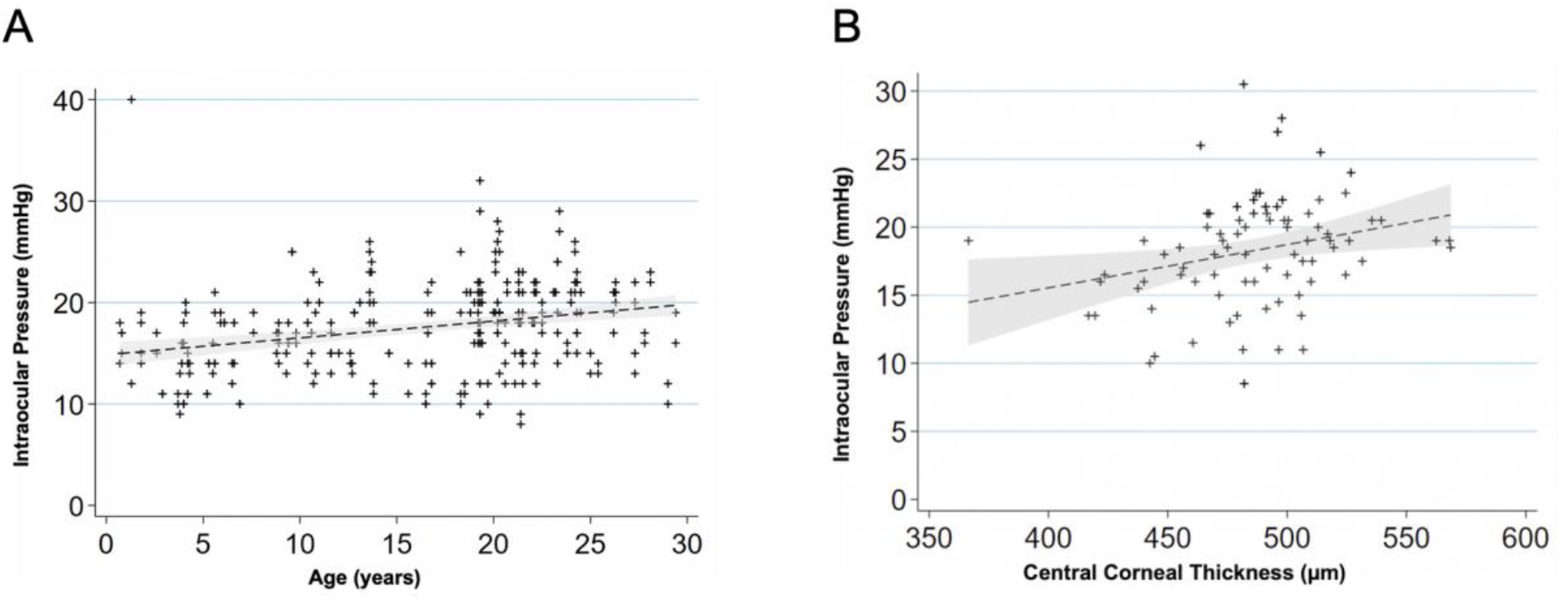
Intraocular Pressure variations with age and central corneal thickness **(A)** Scatterplot showing relationship of age (years) and intraocular pressure (mm Hg) for 142 macaques, 284 eyes. IOP increased by 0.165 mm Hg per increase in year of age (*P* < 0.001). OD (17.3 ± 4.3 mm Hg). OS (17.8 ± 4.7 mm Hg). There was no statistical difference between OD and OS (*P* = 0.369). Males (18.5 ± 4.5 mm Hg). Females (17.1 ± 4.0 mm Hg). There was no statistical difference between males and females (*P* = 0.088). A linear regression line is fitted with a 95% confidence interval in the grey area. **(B)** Scatterplot showing relationship between corneal thickness (μm) and intraocular pressure (mm Hg) (n = 84 primates, 168 eyes). For every 50 μm increase in corneal thickness the intraocular pressure increases 1.5 mm Hg (*P* < 0.001). A linear regression line is fitted with a 95% confidence interval in the grey area. Abbreviation: Mean ± Standard Deviation. OD = right eye, OS = left eye

Central corneal thickness (CCT) measurements were collected from 84 primates, 168 eyes total (486 μm ± 38 μm) with no statistically significant difference between sexes (*P* = 0.479). The IOP and CCT had a significant positive correlation (Figure 2B) with an increase of 1.5 ± 0.4 mm Hg for every 50 μm in CCT (*P* < 0.001). Lens thickness and axial length measurements were collected from 114 individuals, 228 eyes total. The mean lens thickness was 4.24 ± 0.53mm in all animals; 4.20 ± 0.23 mm in males and 4.25 ± 0.50 mm in females (Table 1). There was no significant difference between sexes (*P* = 0.363). Lens thickness measurements suggested a positive correlation of 7.3 ± 5.2 μm per year in age (*P* = 0.162, Figure 3A). The axial length was higher in males (20.06 ± 0.95 mm) than in females (19.67 ± 0.96 mm) when using a generalized linear model to correlate the data from the two eyes of each animal (*P* = 0.046). Axial length increases by 52.82 ± 11.36 μm per year of age (*P* < 0.001, Figure 3B).

**Table 1.** Ocular Biometry

|  | Male: mean<br>(stdev) | Female: mean<br>(stdev) | *P*-value (Males vs<br>Females) |
| --- | --- | --- | --- |
| **Lens Thickness (mm)** | 4.2 (0.2) | 4.3 (0.5) | 0.363 |
| **Axial Length (mm)** | 20.1 (1.0) | 19.7 (1.0) | 0.051 |

Axial length, Lens thickness (n = 114, 228 eyes)

**Figure 3.**
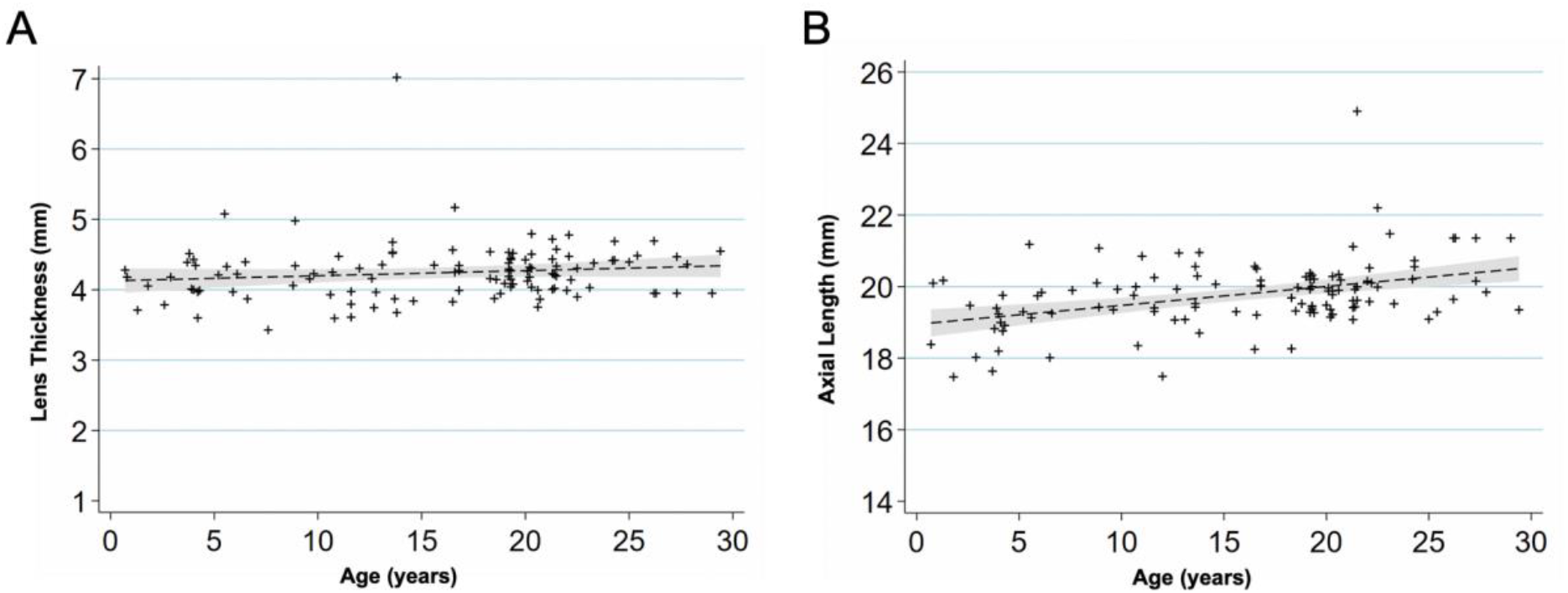
Age-Related Changes in Ocular Biometry **(A)** Scatterplot showing relationship between age (years) and lens thickness (mm) (n = 114 primates, 228 eyes). The lens thickness increases 7.3 μm per increase in year of age (*P* = 0.162). A linear regression line that has been fitted with a 95% confidence interval in the grey area. **(B)** Scatterplot showing relationship between age (years) and axial length (mm) (n = 114 primates, 228 eyes). The axial length increases 52.8 μm per increase in year of age (*P* < 0.001). A linear regression line that has been fitted with a 95% confidence interval in the grey area.

### Retinal Findings and Fundus Measurements

Fundus imaging was performed on 78 individuals; 77 right eyes, 78 left eyes (Table 2). Observations by indirect ophthalmoscopy revealed yellow-white punctate macular lesions of lipoidal degeneration^16–19^ present in 66 individuals (age range = 4.2-29.4 years, mean age = 17.8 years, median age = 19.3 years, 47% of total study subjects). These lesions, reminiscent of small hard drusen in humans, were identifiable on fundus imaging (Figure 4A-D) and were distinct from soft drusen-like macular lesions (Figure 4E) observed in a smaller subset of animals. There were 74 individuals without punctate macular lesions (age range = 0.7-27.8 years, mean age = 14.6 years, median age = 18.8 years). The age distributions of individuals with and without punctate macular lesions are shown in Figure 4F. Forty-two individuals had punctate macular lesions visible by indirect ophthalmoscopy and on fundus images. The mean age of the animals with punctate macular lesions (17.8 ± 6.2 years) was statistically higher than the mean age of animals without them (14.6 ± 8.1 years), suggesting punctate macular lesions are an age-related phenomena (*P* = 0.009).^16–19^ Further grouping of individuals with various degrees (graded as few, moderate, or extensive) of punctate macular lesions is shown in Figure 4G. While the presence of punctate macular lesions may be age-related, the number of punctate macular lesions may not correlate with age. Hematoxylin and eosin stain of a histological section of a rhesus macaque retina clinically graded with extensive punctate macular lesions shows numerous translucent spheroidal lesions in the outer retina in the region of the photoreceptor outer segments and the retinal pigmented epithelium (Figure 4H).

**Table 2.** Fundus measurements (n = 78 primates, 155 eyes)

|  | All<br>Animals:<br>mean<br>(Stdev) | OD:<br>mean<br>(Stdev) | OS:<br>mean<br>(Stdev) | *P*-value<br>(OD vs<br>OS) | Male:<br>mean<br>(Stdev) | Female:<br>mean<br>(Stdev) | *P*-value<br>(Males<br>vs<br>Females) |
| --- | --- | --- | --- | --- | --- | --- | --- |
| Artery (μm<br>diameter) | 50.4<br>(13.8) | 51.8<br>(14.1) | 49.1<br>(13.4) | 0.220 | 53.0<br>(9.4) | 49.3<br>(11.6) | 0.150 |
| Vein (μm<br>diameter) | 125.4<br>(25.1) | 124.4<br>(26.1) | 126.4<br>(24.2) | 0.616 | 132.6<br>(22.9) | 122.7<br>(21.2) | 0.094 |
| Height of Nerve<br>(μm) | 1810.4<br>(143.7) | 1816.7<br>(137.5) | 1804.2<br>(150.8) | 0.590 | 1814.3<br>(144.9) | 1808.2<br>(135.8) | 0.868 |
| Width of Nerve<br>(μm) | 1279.2<br>(127.2) | 1282.5<br>(127.7) | 1275.8<br>(128.2) | 0.745 | 1276.6<br>(148.6) | 1280.6<br>(112.4) | 0.910 |
| Foveal Center to<br>Nerve Center<br>(μm) | 4110.8<br>(287.0) | 4068.0<br>(287.9) | 4153.1<br>(282.4) | 0.065 | 4117.4<br>(258.9) | 4110.6<br>(264.5) | 0.919 |
| Foveal Center to<br>Nerve Edge<br>(μm) | 3441.8<br>(281.2) | 3390.1<br>(279.4) | 3492.9<br>(276.2) | 0.022 | 3445.8<br>(224.5) | 3442.5<br>(261.4) | 0.959 |
| How High Up the Nerve? ($\mu\text{m}$) and %percentage of nerve height | 595.6<br>(218.0)<br>32.9% | 636.1<br>(221.4)<br>35.1% | 555.6<br>(208.4)<br>30.8% | 0.021 | 574.0<br>(155.3),<br>(31.6%) | 600.9<br>(173.8),<br>(33.3%) | 0.516 |
| Area of Avascularization ($\mu\text{m}^2$) | 410.3<br>(153.3) | 405.2<br>(158.9) | 410.7<br>(147.7) | 0.824 | 394.0<br>(132.8) | 413.1<br>(159.9) | 0.453 |
| Area of Avascularization with FA ($\mu\text{m}^2$) | 372.1<br>(140.9) | 345.5<br>(106.5) | 396.8<br>(167.0) | 0.349 | 330.9<br>(128.7) | 400.4<br>(145.9) | 0.205 |

Abbreviation: OD = right eye, OS = left eye, FA = fluorescein angiogram

**Figure 4.**
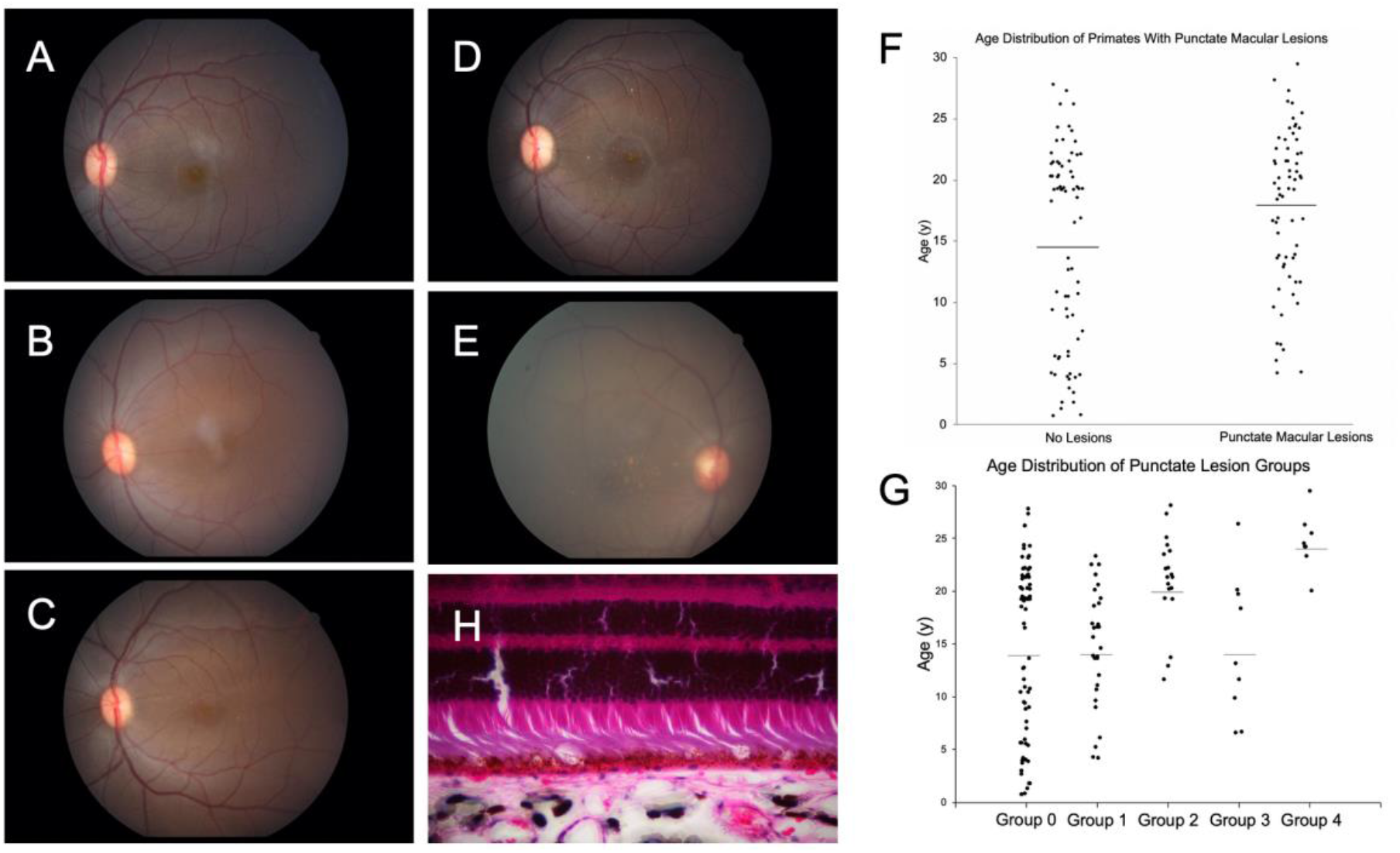
Macular Lesions and Age Distribution **(A)** Color fundus photo Group 0: no punctate macular lesions. **(B)** Color fundus photo Group 1: few punctate macular lesions. **(C)** Color fundus photo Group 2: moderate punctate macular lesions. **(D)** Color fundus photo Group 3: extensive punctate macular lesions. **(E)** Color fundus photo Group 4: soft drusen-like macular lesions. **(F)** Age distribution of rhesus macaques with and without punctate macular lesions (*P* = 0.009). No punctate macular lesions (n = 74, age range = 0.7-27.8 years, mean age = 14.6 years, median age = 18.8 years). Punctate macular lesions (n = 66, age range = 4.2-29.4 years, mean age = 17.8 years, median age = 19.3 years). Macaques with punctate macular lesions = 47% of total study primates. Line represents mean age. **(G)** Age distribution of rhesus macaques with various levels of punctate lesions. Group 0: no punctate lesions (n = 74, age range = 0.7-29.0 years, mean age = 14.6 years, median age = 19.8 years). Group 1: few small hard lesions (n = 30, age range = 4.2-23.3 years, mean age = 14.9 years, median age = 16.0 years). Group 2: moderate small hard lesions (n = 19, age range = 11.6-28.1 years, mean age = 20.9 years, median age = 21.3 years). Group 3: extensive small hard lesions (n = 9, age range = 6.5-26.3 years, mean age = 14.7 years, median age = 13.1 years). Group 4: soft lesions (n = 8, age range = 20.0-29.4 years, mean age = 24.6 years, median age = 24.3 years). Line represents mean age. **(H)** Hematoxylin and eosin stain of a histological section of a rhesus macaque retina clinically graded with extensive punctate macular lesions. Numerous translucent spheroidal lesions in the outer retina in the region of the photoreceptor outer segments and the retinal pigmented epithelium are seen. The section is from the macular region as evidenced by multiple rows of nuclei in the retinal ganglion cell layer.

**Figure 5.**
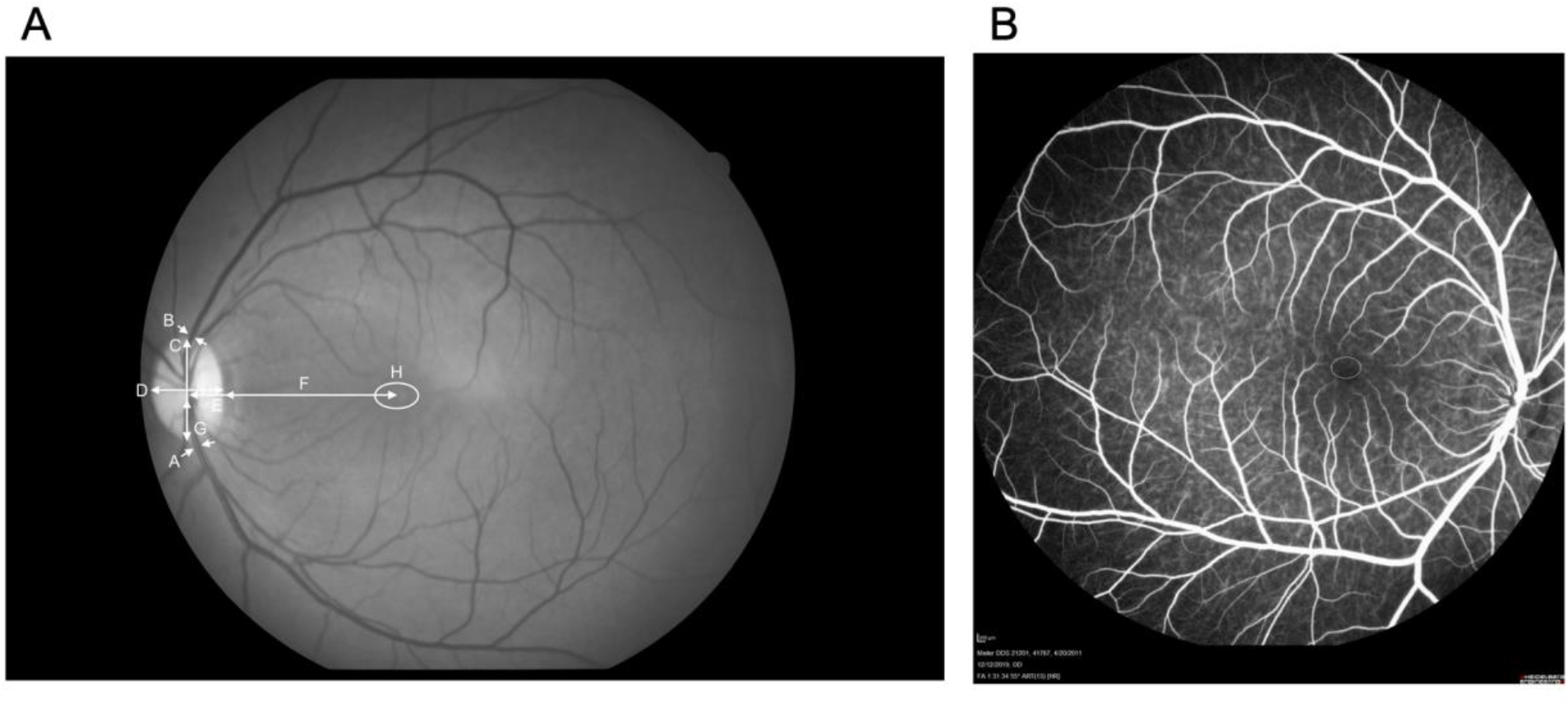
Fundus Images and Measurements (n = 78 primates, 155 eyes) **(A)** Red-free fundus photo with measurements. The arrows represent measurements taken. A: artery (diameter). B: vein (diameter). C: height of optic nerve. D: width of optic nerve. E: foveal center to nerve center. F: foveal center to nerve edge. G: relative position of the foveal center from base of the optic nerve. H: area of avascularization. **(B)** Fluorescein angiogram showing FAZ measurement.

Fundus measurements for the 151 eyes with photographs are summarized in Table 2. The optic nerve had a maximal vertical diameter of 1.8 mm while the horizontal diameter was 1.3 mm. The disc-fovea distance in rhesus monkeys was 4.1 ± 0.3 mm. The fovea of the rhesus macaque sits 32.9% up the optic nerve vertical diameter. This results in a disc-fovea angle of approximately 8.2°. The diameter of the main branches of the central retinal artery and vein at the point of crossing the edge of the optic disc are 50.5 μm and 125.4 μm, respectively. The arterial-venous ratio (AVR) was calculated to be 0.43 ± 0.14. The apparent foveal avascular zone (FAZ) using fundus photography was 0.4 ± 0.2 mm^2^. The ratio of the vertical diameter (0.6 mm) to horizontal diameter (0.9 mm) was 0.70 ± 0.19. When measuring FAZ area with fluorescein angiography, similar values were found, 0.4 ± 0.1 mm^2^ (n =17, eyes = 27). The vertical diameter was 0.6 mm and the horizontal diameter was 0.8 mm. The vertical to horizontal ratio using fluorescein angiography was 0.68 ± 0.18. For primates that had both fundus imaging and fluorescein angiography images (n = 9, eyes = 11) the FAZ measured 0.5 ± 0.1 mm^2^ and 0.3 ± 0.1 mm^2^ respectively.

Some differences between left and right eyes were identified. The distance from the foveal center to the optic nerve edge between the OD and the OS measured 3.4 ± 0.3 mm in the OD and 3.5 ± 0.3 mm in the OS (*P* = 0.022). The relative distance of the foveal center up the optic nerve vertical height measured 0.6 ± 0.2 mm (35.1%) the OD and 0.6 ± 0.2 mm (30.8%) the OS (*P* = 0.021). The biological relevance of this left-right axis asymmetry is unclear. There were no significant differences between male and female fundus measurements.

### Electroretinography (ERG)

Full-field ERG measurements were collected from 75 individuals, 150 eyes total. The average amplitudes and latencies from dark-adapted and light-adapted tests are shown in Table 3. The full field ERG algorithm resulted in a-wave (chiefly representing photoreceptor activity) and b-wave (largely representing bipolar cell activity) times and amplitudes. The dark-adapted dim stimulus test (0.01) represents rod pathway function. The dark-adapted bright flash tests (3.0 and 10.0) correspond to the combined responses of the rod and cone pathways. The cone pathway is the main contributor to the light adapted tests (3.0 and flicker). Oscillatory potentials (OP), a representation of extracellular electrical currents between bipolar, amacrine, and ganglion cells in the inner plexiform layer^20–22^, were also calculated and presented.

**Table 3.** Full-field electroretinogram measurements

| | | Latency (ms) | | Amplitude ($\mu\text{V}$) | |
| --- | --- | --- | --- | --- | --- |
|  |  | Mean | Stdev | Mean | Stdev |
| Dark 0.01 | b-wave | 87.9 | 14.2 | 68.8 | 45.2 |
| Dark 3.0 | a-wave | 18.2 | 2.3 | -90.9 | 37.0 |
|  | b-wave | 45.1 | 5.3 | 191.6 | 73.2 |
| Dark 10.0 | a-wave | 15.6 | 2.2 | -122.5 | 47.2 |
|  | b-wave | 45.4 | 6.0 | 207.2 | 80.1 |
| Light 3.0 | a-wave | 15.0 | 1.2 | -16.4 | 7.6 |
|  | b-wave | 28.9 | 1.7 | 68.3 | 29.9 |
| Flicker | amplitude | 25.5 | 1.4 | 51.0 | 22.2 |
| OP1 | amplitude | 18.3 | 2.0 | 14.8 | 10.9 |
| OP2 | amplitude | 24.9 | 2.3 | 12.4 | 8.4 |
| OP3 | amplitude | 31.7 | 2.7 | 16.3 | 11.1 |
| OP4 | amplitude | 39.4 | 2.8 | 11.5 | 7.3 |
| OP5 | amplitude | 46.0 | 2.8 | 5.5 | 4.1 |
| OP sum | amplitude | 145.3 | 15.2 | 57.8 | 34.5 |

Abbreviation: OP = oscillatory potential
Dark 0.01 (n = 73, eyes = 146)
Dark 3.0, Dark 10.0, Flicker, OPs 1-4 and sum (n = 75, eyes = 150)
Light 3.0 (n = 71, eyes = 142)
OP5 (n = 65, eyes = 130)

### Optical Coherence Tomography Segmentation

Tomographic images of the macula were obtained from 60 individuals with 49 right eyes and 49 left eyes (Table 4). To determine if there are structural changes in the retina over time, individual retinal layer thicknesses were plotted against age (Figure 6). Significant changes in layer thickness with increase in age was found in the retinal ganglion cell layer (GCL), inner nuclear layer (INL) temporal to the foveal center, photoreceptor outer segments (OS), choriocapillaris (CC), and outer choroid (OC). Significant reduction in thickness was found in the GCL both nasal (*P* = 0.019) and temporal (*P* = 0.001) to the fovea. The nasal side decreased by 0.30 μm and the temporal side by 0.37 μm for each increase in year of age. The thickness of the INL reduced only temporal to the foveal by 0.19 μm per increase in year of age (*P* = 0.025). The OS increased in thickness on the nasal side (0.16 μm per year, *P* = 0.003), temporal side (0.26 μm per year, *P* < 0.001), and foveal center (0.36 μm per year, *P* < 0.001). The CC thickness increased in the nasal side (0.34 μm per year, *P* < 0.001), temporal side (0.38 μm per year, *P* < 0.001), and foveal center (0.27 μm per year, *P* < 0.001). Finally, the OC thickness increased in the nasal side (3.66 μm per year, *P* < 0.001), temporal side (3.20 μm per year, *P* < 0.001), and foveal center (3.58 μm per year, *P* < 0.001). The other layers were not found to vary significantly with age.

**Table 4.** Retinal structural measurements from optical coherence tomography (n = 60 primates, 98 eyes)

|  | Nasal<br>(mean) | Nasal<br>(Stdev) | Temporal<br>(mean) | Temporal<br>(Stdev) | Fovea<br>(mean) | Fovea<br>(Stdev) |
| --- | --- | --- | --- | --- | --- | --- |
| NFL<br>($\mu\text{m}$) | 21.4 | 5.8 | 15.4 | 3.1 | | |
| GCL<br>($\mu\text{m}$) | 39.4 | 8.7 | 33.3 | 8.9 | 16.0 | 4.7 |
| IPL<br>($\mu\text{m}$) | 49.6 | 7.3 | 47.8 | 6.6 | | |
| INL<br>($\mu\text{m}$) | 44.0 | 5.4 | 41.6 | 6.0 | | |
| OPL<br>($\mu\text{m}$) | 26.9 | 4.0 | 26.9 | 5.6 | | |
| ONL<br>($\mu\text{m}$) | 63.0 | 7.5 | 61.1 | 7.2 | 82.2 | 15.7 |
| IS (μm) | 35.4 | 3.5 | 34.9 | 4.1 | 39.5 | 4.4 |
| OS (μm) | 23.1 | 4.1 | 23.8 | 5.1 | 37.6 | 6.7 |
| RPE (μm) | 33.8 | 5.2 | 34.6 | 5.7 | 29.7 | 5.2 |
| CC (μm) | 10.9 | 4.2 | 12.1 | 4.7 | 11.8 | 4.2 |
| OC (μm) | 194.3 | 65.8 | 198.4 | 62.8 | 200.9 | 60.6 |
| TRT (μm) | 336.6 | 17.6 | 319.4 | 17.2 | 205.1 | 18.8 |

Abbreviations: CC: choriocapillaris, GCL: ganglion cell layer, INL: inner nuclear layer, IPL: inner plexiform layer, IS: inner segments of photoreceptors, NFL: nerve fiber layer, OC: outer choroid, ONL: outer nuclear layer, OPL: outer plexiform layer, OS: outer segments of photoreceptors, RPE: retinal pigment epithelium. TRT: total retinal thickness.

**Figure 6.**
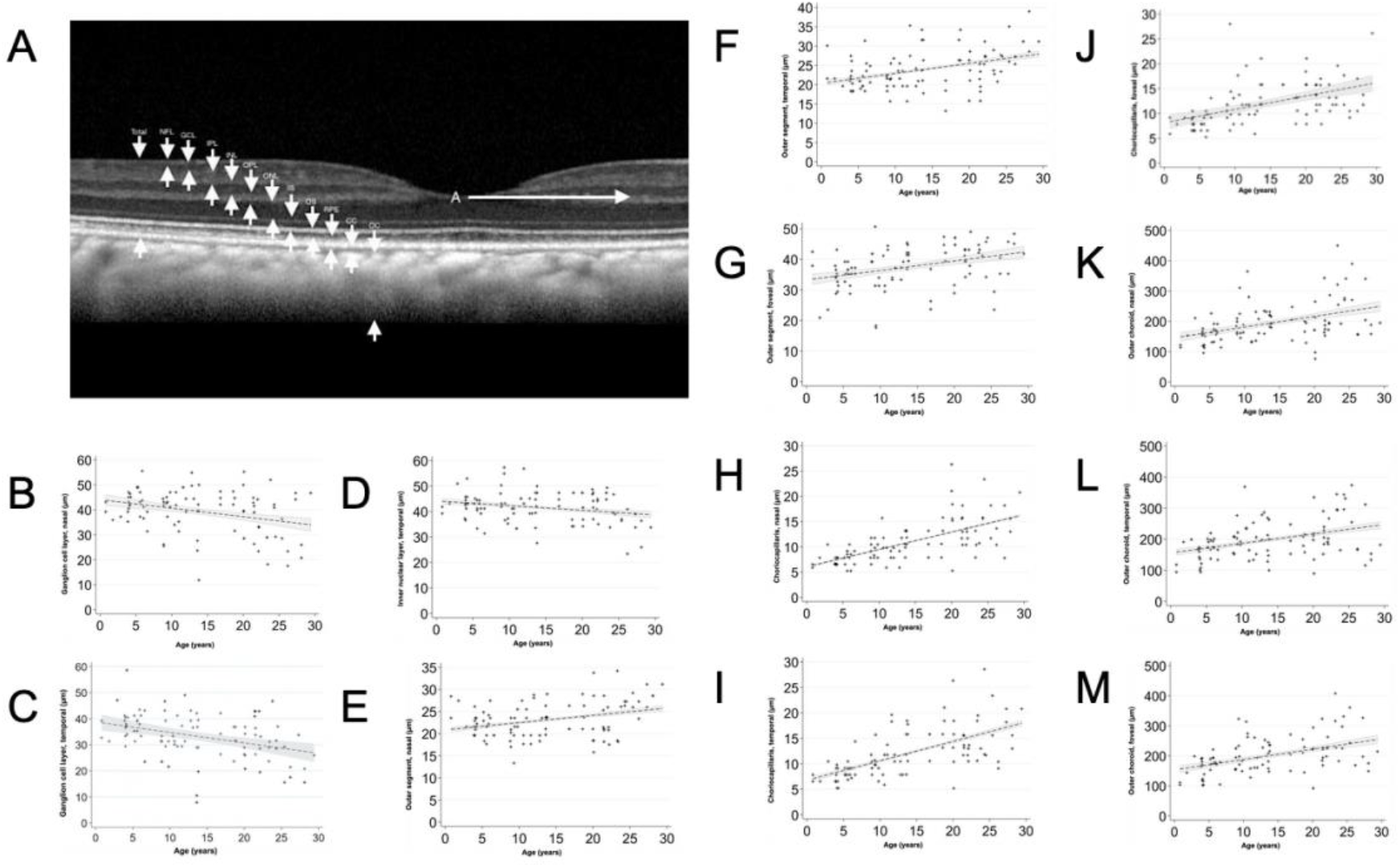
Optical Coherence Tomography (OCT) Images and Measurements **(A**) Measurement of retinal and choroidal layers (n = 60 primates, 98 eyes). White arrow A: 1000μm from foveal center. All retinal layers were measured at this distance on both temporal and nasal sides. Total retinal thickness is measured from the nerve fiber layer (NFL) to the retinal pigment epithelium (RPE). **(B)** Scatterplot showing relationship between age (years) and GCL nasal side (μm). GCL decreased by a factor of 0.30 μm per increase in year of age (*P* = 0.019). **(C)** Scatterplot showing relationship between age (years) and GCL temporal side (μm). GCL decreased by a factor of 0.37 μm per increase in year of age (*P* = 0.001). **(D)** Scatterplot showing relationship between age (years) and INL temporal side (μm). INL decreased by a factor of 0.19 μm per increase in year of age (*P* = 0.025). **(E)** Scatterplot showing relationship between age (years) and OS nasal side (μm). OS increased by a factor of 0.16 μm per increase in year of age (*P* = 0.003). **(F)** Scatterplot showing relationship between age (years) and OS temporal side (μm). OS increased by a factor of 0.26 μm per increase in year of age (*P* < 0.001). **(G)** Scatterplot showing relationship between age (years) and OS foveal side (μm). OS increased by a factor of 0.36 μm per increase in year of age (*P* < 0.001). **(H)** Scatterplot showing relationship between age (years) and CC nasal side (μm). CC increased by a factor of 0.34 μm per increase in year of age (*P* < 0.001). **(I)** Scatterplot showing relationship between age (years) and CC temporal side (μm). CC increased by a factor of 0.38 μm per increase in year of age (*P* < 0.001). **(J)** Scatterplot showing relationship between age (years) and CC foveal side (μm). CC increased by a factor of 0.27 μm per increase in year of age (*P* < 0.001). **(K)** Scatterplot showing relationship between age (years) and OC nasal side (μm). CC increased by a factor of 3.66 μm per increase in year of age (*P* < 0.001). **(L)** Scatterplot showing relationship between age (years) and OC temporal side (μm). CC increased by a factor of 3.20 μm per increase in year of age (*P* < 0.001). **(M)** Scatterplot showing relationship between age (years) and OC nasal side (μm). CC increased by a factor of 3.58 μm per increase in year of age (*P* < 0.001). Each scatterplot has a linear regression line that has been fitted with a 95% confidence interval in the grey area. Abbreviations: NFL: nerve fiber layer. GCL: ganglion cell layer. IPL: inner plexiform layer. INL: inner nuclear layer. OPL: outer plexiform layer. ONL: outer nuclear layer. IS: inner segments. OS: outer segments. RPE: retinal pigmented epithelium. CC: choriocapillaris. OC: outer choroid. TRT: total retinal thickness.

To determine if the retinal nerve fiber layer and ganglion cell layers measured on the macular OCT have a relationship with intraocular pressure measured in the eye, we plotted the combined NFL+GCL thickness of each eye against the IOP measured in that eye (Figure 7). A linear regression was performed after correlating the eyes of each individual animal, which revealed the thickness of these layers decreased both nasal (Figure 7A, 0.6 μm for each 1 mm Hg increased IOP, *P* = 0.035) and temporal (Figure 7B, 0.9 μm for each 1 mm Hg increased IOP, *P* = 0.001) to the foveal center.

**Figure 7.**
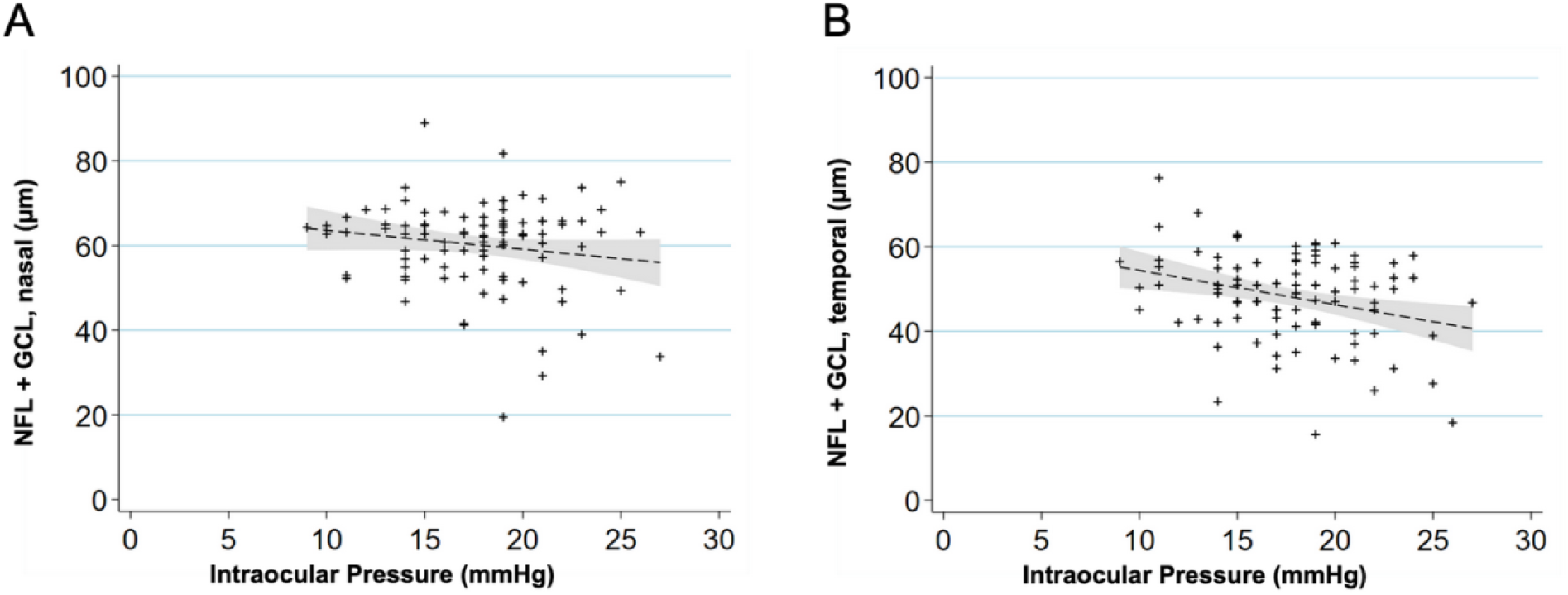
Relationship between intraocular pressure and retinal nerve fiber layer and ganglion cell layer. **(A)** Scatterplot showing relationship between IOP (mm Hg) and NFL+GCL nasal side (μm). NFL+GCL decreased by a factor of 0.6 μm per 1 mm Hg increase (*P* = 0.035). A linear regression line that has been fitted with a 95% confidence interval in the grey area. **(B)** Scatterplot showing relationship between IOP (mm Hg) and NFL+GCL temporal side (μm). NFL+GCL decreased by a factor of 0.9 μm per 1 mm Hg increase (*P* = 0.001). A linear regression line that has been fitted with a 95% confidence interval in the grey area.

### Quantitative Autofluorescence (qAF) analysis

The qAF8 data were included from 68 eyes of 44 individuals, while 57 eyes from 38 individuals were discarded for poor quality (54.4% inclusion rate). A typical en-face image in Heidelberg’s qAF mode is shown in Figure 8A. The values of the 8 segments of the middle ring of the pattern (qAF8) were analyzed. The mean qAF8 value for all individuals was 91.4 autofluorescence units (standard deviation 31.6). We observed an increase in qAF8 by a factor of 1.021 units per increase in year of age (*P* = 0.006, Figure 8B). No difference was observed in qAF8 between male and female sexes (*P* = 0.102).

**Figure 8.**
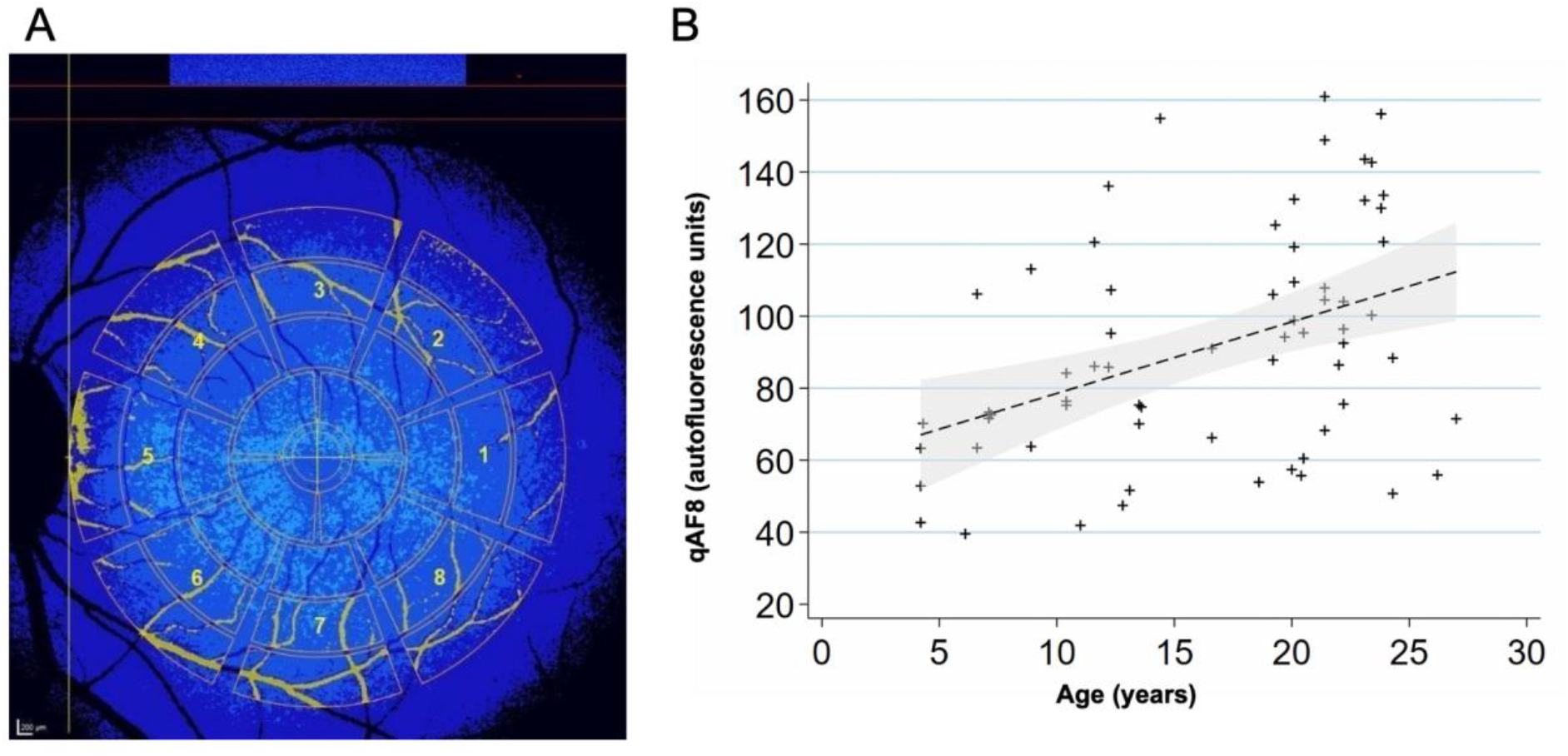
Quantitative autofluorescence analysis (n = 44 primates, 68 eyes) **(A)** To acquire qAF8, the eight middle segments (numbered octants) of the perifoveal Delori grid was used in Heidelberg’s qAF mode. The center circle is placed over the foveal center and the grid is expanded until it touches the tangential edge of the optic nerve. Vessels are automatically excluded and the qAF measurements in each octant are normalized to the internal autofluorescence standard shown in the blue bar at the top of the image. **(B)** Scatterplot showing relationship between qAF8 and age (years). Mean (standard deviation): 91.4 (31.3). The mean qAF8 increased by a factor of 1.021 autofluorescence units per increase in year of age (*P* = 0.006). A linear regression line has been fitted with a 95% confidence interval in the grey area.

## Discussion

### Structural Similarities to Human Eyes: Intraocular Pressure (IOP), Central Corneal Thickness (CCT), Lens Thickness, Axial Length, and Fundus Structural Measurements

The eye of the rhesus macaque shares many anatomical similarities to the human eye. Mean IOP was 18 ± 4 mm Hg and thus comparable to normal human IOP values which have classically been considered to range between 10 and 21 mm Hg.^23,24^ Hollows and colleagues examined 4,231 people and found women had an approximate increase of 1 mm Hg compared with men. They also observed a significant increase in IOP with age. More recent studies in a large Chinese population, 3135 normal patients, have demonstrated a similar mean,^25^ suggesting little variation across human ethnicities. A comparison between rhesus macaques and humans is represented in Table 5. In our cohort of rhesus macaques, we also observed a statistically significant increase in IOP with age (*P* < 0.005) but did not find a significant difference between males and females.

**Table 5.** Comparison of mean intraocular pressure.

|  | Rhesus macaque | Human<sup>24</sup> | Human<sup>25</sup> |
| --- | --- | --- | --- |
| Male (mm Hg) | 18.5 ± 4.5 | 15.9 ± 3 |  |
| Female (mm Hg) | 17.1 ± 4.0 | 16.6 ± 3 |  |
| Overall (mm Hg) | 17.6 ± 4.2 |  | 14.7 ± 2.8 |

Abbreviations: Mean ± Standard Deviation.

Central corneal thickness (CCT) was determined to be 486.0 ± 37.6 μm in our cohort of rhesus macaques, and is thus mildly thinner to that reported in humans 535 ± 31 μm.^26^ In that meta-analysis, there was a positive correlation of 1.1 ± 0.6 mm Hg in IOP change for each 10% change (~50 μm) in CCT. We found a similar correlation between IOP and corneal thickness of approximately 1.5 ± 0.4 mm Hg for every 50μm in corneal thickness (*P* < 0.001).

The fully developed normal human lens is 4.0 mm in thickness at age 20, but increases to 4.3 mm at the age 40 of years, 4.45 mm at 50 years, 4.7 mm at 60 years, and above 4.7 mm after 60 years of age.^27^ The rate of increase in crystalline lens thickness has been estimated at 0.15–0.20 mm per decade of life.^28^ The rhesus macaque lens is 4.24 ± 0.41 mm in thickness. A previous study did find a significant age-related increase of 0.03 ± 0.01 mm per year (*P* = 0.002) when lens thickness was studied within a smaller cohort of rhesus monkeys.^29^ However, we did not find a significant age-related increase in lens thickness in our animals (*P* = 0.162). Axial length measurement of the rhesus eye was determined to be 19.77 ± 0.97 mm with a significant age-related increase. In humans, the newborn eye is 16 mm and continues to grow to approximately 24 mm.^30^ More precise measurement of normal emmetropic human eyes revealed axial length measurements of about 23.3 ± 0.5 mm with broader variation among ametropic individuals.^31^ Therefore, the rhesus macaque eye is ~85% the size of the human eye.

Structural measurements between the human and rhesus fundi were similar. The optic nerve in the rhesus macaque had a vertical diameter of 1.8 mm while the horizontal diameter was 1.3 mm resulting in a vertical to horizontal ratio of 1.42 ± 0.10. In a study by Jonas and Hayreh, they also documented optic disc shape in seventeen rhesus macaques and obtained a similar vertical to horizonal disc diameter of 1.39 ± 0.09.^32^ In the human eye, the optic disc is more circular with a vertical diameter of 1.88 mm and a horizontal diameter 1.77 mm.^33^ The disc-fovea distance is consistent between adult humans and has been reported as 4.78 ± 0.06 mm.^34^ This has been confirmed in a large study of 2836 eyes and was reported as 4.76 ± 0.34 mm.^35^ The disc-fovea distance in rhesus monkeys was 4.1 ± 0.3 mm, similar to but less than humans, consistent with the slightly (~14%) smaller rhesus eye. Mok and Lee found the human optic nerve head area was 2.79 mm^2,34^ and we report a value of 2.3 mm^2^ revealing that the rhesus macaque optic disc area is approximately 83% of its human counterparts. Notably, the rhesus eye measures 85% of the human eye in axial length and 86% of the human eye in disc-fovea distance, suggesting very similar interspecies structural ocular anatomic proportionality.

The disc-fovea angle in a large cohort of 3052 people was reported as 7.76 ± 3.63°.^36^ This results in the fovea lying 0.65 mm upward from the inferior edge of the optic disc, approximately 34.6% (0.65/1.88 mm) of the way to the top of the optic disc height. The fovea of the rhesus macaque eye sits 32.9% up the optic nerve height. This results in disc-fovea angle of approximately 8.2°. In humans, the thickness of the main branches of the central retinal artery and vein at the point of crossing the edge of the optic disc have been reported as 169.8 μm and 242.1 μm, respectively.^37^ The corresponding values from the rhesus macaque retina are 50.5 μm for arterioles and 125.4 μm for venules. Our reported AVR for males (0.42 ± 0.11) and females (0.41 ± 0.13) are both thinner than that reported in both male (0.87 ± 0.08) and female (0.89 ± 0.08) humans.^38^ The foveal avascular zone (FAZ) is 0.27 mm^2^ in healthy human volunteers using optical coherence tomography angiography (OCTA), which reveals the retinal microvasculature with very high resolutions.^39^ Angiographic imaging of the foveal avascular zone in rhesus macaques was not available for this study. In the macaque, the FAZ was 0.4 ± 0.2 mm^2^ with a vertical diameter of 0.6 mm and a horizontal diameter of 0.9 mm. This may be an overestimation since these measurements were based on fundus photography, a modality incapable of showing the microvasculature at the innermost FAZ boundary. By using fluorescein angiography, a more accurate representation of the FAZ was visible which corresponded to 0.4 ± 0.1 mm^2^ with a vertical diameter of 0.6 mm and a horizontal diameter of 0.8 mm. When comparing FAZ areas within the same primates, the difference between imaging modalities is more clearly seen. The FAZ using fundus imaging was 0.5 ± 0.1 mm^2^ and using fluorescein angiography was 0.3 ± 0.1 mm^2^. Both modalities show the macaque FAZ is ovoid in shape in contrast to the more circular FAZ seen in humans.

### ERG Similarities to Human Eyes

Electroretinographic recordings in rhesus macaque are similar to humans and other NHPs. A study by Bouskila et al^40^ standardized ERG measurements in the Green Monkey (*Chlorocebus sabaeus*) in which they obtained similar latency results as those reported here (Table 6). Our closest amplitude results were seen in maximal cone and rod response (dark 3.0) b-wave. In comparison to human data, the primate retina has an overall minimally increased latency as compared to the human, but amplitudes appeared similar. Liu et al^41^ determined a mean latency for dark 0.01 b-wave (83 ms) and dark 3.0 a-wave (15 ms) and b-wave (44 ms). Our results for each of these tests were 87.9ms, 18.2ms, and 45.1ms, respectively. These latency results correlate to humans ages 40-70 years old. Overall, NHP waveforms are very similar to human ERGs. There is a precedent for gender differences in multifocal ERG recordings in rhesus macaque, and also between sexes in cynomolgus macaques.^42^ In this study, we did not report multifocal ERG data. Furthermore, we did not observe differences between female and male rhesus macaques using our full field ERG instrument.

**Table 6.** Comparison of mean electroretinogram values.

|  |  | Rhesus Macaque |  | Green Monkey<sup>40</sup> |  | Human<sup>41</sup> |  |
| --- | --- | --- | --- | --- | --- | --- | --- |
|  |  | Latency (ms) | Amplitude (μV) | Latency (ms) | Amplitude (μV) | Latency (ms) | Amplitude (μV) |
| Dark<br>0.01 | b-wave | 87.9 | 68.8 | 79.9 | 88.9 | 83 | 41 |
| Dark<br>3.0 | a-wave | 18.2 | -90.9 | 14.8 | 115.1 | 15 | 39 |
|  | b-wave | 45.1 | 191.6 | 36.7 | 203.7 | 44 | 61 |
| Dark<br>10.0 | a-wave | 15.6 | -122.5 | 9.8 | 174.9 | 11 | 44 |
|  | b-wave | 45.4 | 207.2 | 30.8 | 230.7 | 47 | 64 |
| Light<br>3.0 | a-wave | 15.0 | -16.4 | 12.3 | 22.1 | 11 | 6 |
|  | b-wave | 28.9 | 68.3 | 27.7 | 81.5 | 28 | 26 |
| Flicker | amplitude | 25.5 | 51.0 | 24.3 | 88.9 | 25 | 29 |
| OP1 | amplitude | 18.3 | 14.8 |  |  |  |  |
| OP2 | amplitude | 24.9 | 12.4 |  |  |  |  |
| OP3 | amplitude | 31.7 | 16.3 |  |  |  |  |
| OP4 | amplitude | 39.4 | 11.5 |  |  |  |  |
| OP5 | amplitude | 46.0 | 5.5 |  |  |  |  |
| OP sum | amplitude | 145.3 | 57.8 | 20.3 | 60.2 |  |  |

### Retinal OCT and qAF Similarities to Human Eyes

The functional and structural characterization of a macular lesion in a rhesus macaque has been reported using OCT, ERG, and histology.^43^ Macular structure is very similar between humans and NHPs, as both have similar retinal lamination and a foveal depression and architecture. Tomagraphic images were measured manually using the ImageJ software. Yiu et al^44^ performed measurements on six rhesus macaques (older than 6 years) using semi-automated segmentation of OCT images (comparison shown in Table 7). Overall, our inner retinal layers thicknesses measured closely to theirs with the exception of the ONL and OS. These differences may be due to the focal measurements we took at 1.5 mm nasal and temporal to the foveal pit rather than across the entire central macular b-scan or to subtle differences between landmarks used at the outer segment-RPE boundary. Our study also compared a larger number of NHP eyes across a broader spectrum of ages. It has been shown that the outer retinal layers continue to develop with age until adulthood is reached.^45,46^ We found a similar trend within our cohort of individuals (Figure 6). Since rhesus macaques generally reach adulthood between 5-20 years old,^47^ our generally older cohort of macaques may have resulted in small variations in outer retinal layers. Some differences may have arisen as a result of semi-automated computer software compared to manual measurements used in this study.

**Table 7.** Comparison of retinal structural measurements using optical coherence tomography.

|  | Parafoveal (RM) | Parafoveal<sup>44</sup> (RM) | Parafoveal<sup>49</sup> (CM) | Parafoveal<sup>48</sup> (H) | Foveal (RM) | Foveal<sup>44</sup> (RM) | Foveal<sup>49</sup> (CM*) | Foveal<sup>48</sup> (H) |
| --- | --- | --- | --- | --- | --- | --- | --- | --- |
| NFL<br>($\mu\text{m}$) | $18 \pm 4$ | $20.6 \pm 2.9$ | | 34 | | $9.2 \pm 2.3$ | | $4 \pm 3$ |
| GCL<br>($\mu\text{m}$) | $36 \pm 9$ | $34.5 \pm 4.1$ | 72 | 89 | $16 \pm 5$ | $12.1 \pm 3.7$ | 25 | $56 \pm 5$ |
| IPL<br>($\mu\text{m}$) | $49 \pm 7$ | $45.3 \pm 4.9$ | 62 | | | $10.6 \pm 5.1$ | 45 | |
| INL<br>($\mu\text{m}$) | $43 \pm 6$ | $39.8 \pm 4.9$ | 77 | 38 | | $12.5 \pm 4.6$ | 50 | $23.3 \pm 3$ |
| OPL<br>($\mu\text{m}$) | $27 \pm 5$ | $27.1 \pm 2.9$ | 62 | 39 | | $13.5 \pm 5.9$ | 175 | $38 \pm 7$ |
| ONL<br>($\mu\text{m}$) | $62 \pm 7$ | $77.6 \pm 8.5$ | 48 | 81 | $82 \pm 16$ | $96.5 \pm 15.3$ | 50 | $102 \pm 7$ |
| IS ($\mu\text{m}$) | $35 \pm 4$ | $32.5 \pm 1.4$ | 48 | | $40 \pm 4$ | $34.0 \pm 1.9$ | 35 | |
| OS<br>($\mu\text{m}$) | $24 \pm 5$ | $39.4 \pm 2.6$ | 29 | 37 | $38 \pm 7$ | $47.0 \pm 3.2$ | 50 | $42 \pm 5$ |
| RPE<br>($\mu\text{m}$) | $34 \pm 6$ | $15.5 \pm 2.1$ | 38 | | $30 \pm 5$ | $14.7 \pm 2.8$ | 55 | |
| CC<br>($\mu\text{m}$) | $12 \pm 5$ | $24.0 \pm 4.3$ | | | $12 \pm 4$ | $23.8 \pm 5.5$ | | |
| OC<br>($\mu\text{m}$) | 196 ±<br>64 | 171.0 ±<br>21.3 | | | 201 ±<br>61 | 174.1 ±<br>18.2 | | |

Abbreviations: Mean ± Standard Deviation. RM = rhesus macaque. CM = cynomolgus monkey. H = human. \* measurements were taken from closest measurement to foveal center.

Consistent with human studies, OCT layers measured slightly thicker on the nasal side compared to the temporal side due to the thicker NFL on the nasal aspect of the macula.^48^ Variations in the definitions of the layers and also measurement of combinations of layers (e.g. IPL+GCL) make accurate comparisons across these studies difficult. While retinal lamination is grossly very similar between human and NHP eyes, some subtle species differences may exist. OCT imaging of the cynomolgus monkey (*Macaca fasciculari*s) has reported proportionally thicker GCL, IPL, INL, OPL, IS retinal layers and a thinner ONL layer than both rhesus macaque and human studies (Table 7).^49^ These small anatomical differences between monkey species support the use of the rhesus macaque as a more precise model for the human retina.

Autofluorescence imaging has been utilized to reveal structural detail within the macaque eye.^50,51^ In rhesus macaques, qAF was significantly lower than in humans with our average of 91.4 qAF units. Greenberg et al^15^ reported a cohort of age 5-60 year old healthy individuals with a calculated average of 283.9 qAF units, while Wang et al^52^ reported a cohort of age 18-78 year-old healthy individuals with 253.6 qAF units. Previous studies in rhesus macaques have demonstrated elevated qAF in animals deficient in macular xanthophylls, such as lutein and zeaxanthin.^53^ Rhesus macaques also have darker uveal pigmentation than humans due to increased melanin from choroidal melanocytes, although the difference in RPE melanin content is unclear.^54^ Melanin is known to contribute to the fundus autofluorescence.^55^ The increased choroidal pigmentation in rhesus monkeys may explain in part the qAF difference between rhesus and human. The limitations of cSLO imaging in rhesus macaques under sedation may also contribute to lower qAF values in our cohort. Based on image quality criteria defined in the manufacturer’s Quantitative Autofluorescence Analysis Software User Manual, even our highest quality images may have lacked brightness, image contrast, and clear visibility of anatomical structures. In order to pick the highest quality images, we discarded nearly half of available primate eyes as they were not of sufficient quality. In spite of the reduced qAF values in rhesus compared to humans, we observed an increase in qAF with increasing age consistent with human data.^15,52,56^ However, despite our inclusion of macaques of all ages, we did not observe a significant decline in qAF units, which occurs at age 75 years in humans.^57^ We did not find a statistically significant difference between males and females (*P* = 0.104), although it has been shown in human data that males have slightly lower qAF than females.^15,52^

Overall, functional, structural, and relevant species adaptations such as the macula appear similar between the rhesus macaque and the human eye. Our ability to compare normative data across a large cohort of macaques allowed us to make measurements and comparisons with relative precision. The summary of all collected ocular biometric data from our study show that the rhesus macaque eye is proportional to and highly similar in every measurable aspect to the human eye, while being roughly 15% smaller. Age-related changes documented in humans such as IOP elevation, axial length increase, presence of drusen-like punctate macular lesions of lipoidal degeneration, and increasing qAF were confirmed in the eye of rhesus macaques. The positive correlation between CCT and measured IOP was also confirmed. Furthermore, we have shown that IOP correlates with thinner NFL+GCL layer on OCT, prompting future investigation into the degree to which primary open angle glaucoma is recapitulated in rhesus macaques. Our findings support the use of the NHP eye as a model for advanced translational vision science research, especially those related to macular and cone-disorders and age-related diseases.

## Acknowledgements

The authors thank Monica Motta and Michelle Ferneding for expertise in ophthalmic imaging, electrophysiology, data management, and research support.

## Notes

Conflicts of interest: The authors declare no conflicts of interest related to this study.

Financial support: Ala Moshiri is supported by NIH K08 EY027463, NIH U24 EY029904, and Barr Foundation for Retinal Research. Sara M. Thomasy is supported by NIH R01 EY016134, and NIH U24 EY029904. Timothy Stout, Rui Chen, and Jeffrey Rogers are supported by NIH U24 EY029904. This research was also supported by an unrestricted grant from Research to Prevent Blindness to Baylor College of Medicine. Glenn C. Yiu is supported by NIH K08 EY026101, NIH R21 EY031108, the Brightfocus Foundation, and Macula Society. Tu M. Tran was supported by Fight for Sight SS-19-001. No funding organizations had any role in the design or conduct of this research. The content is solely the responsibility of the authors and does not necessarily represent the official views of the funding agencies.

### Competing Interest Statement

The authors have declared no competing interest.

